## Supplemental Figures for "N-cadherin dynamically regulates pediatric glioma cell migration in complex environments"

**Figure S1** (related to Fig. 1) N-cad expression, depletion, and controls for PHGG migration assays.
**Figure S2** (related to Fig. 3 and 4) Role of intercellular N-cad homotypic interactions and importance of N-cad for catenin localization.
**Figure S3** (related to Fig. 5) N-cad endocytosis, recycling and surface levels in leader and follower cells.
**Figure S4** (related to Fig. 6) YAP1 signaling and wound healing gene expression is increased in leader cells.
**Figure S5** (related to Fig. 8) YAP1/TAZ regulates PHGG migration and N-cad endocytosis.
**Table S1** (related to Fig. 1) mRNA expression levels of cell adhesion receptors in patient-derived PHGGs.
**Table S2** (related to Fig. 1) RNA sequencing results comparing control and N-cad shRNA cells.
**Table S3** (related to Fig. 6) RNA sequencing results comparing leader and follower cells.
**Video 1** (related to Fig. 1) Control and N-cad shRNA spheroid migration on laminin.
**Video 2** (related to Fig. 6) Migrating leader and follower cells on neurons or laminin.

### Figure S1

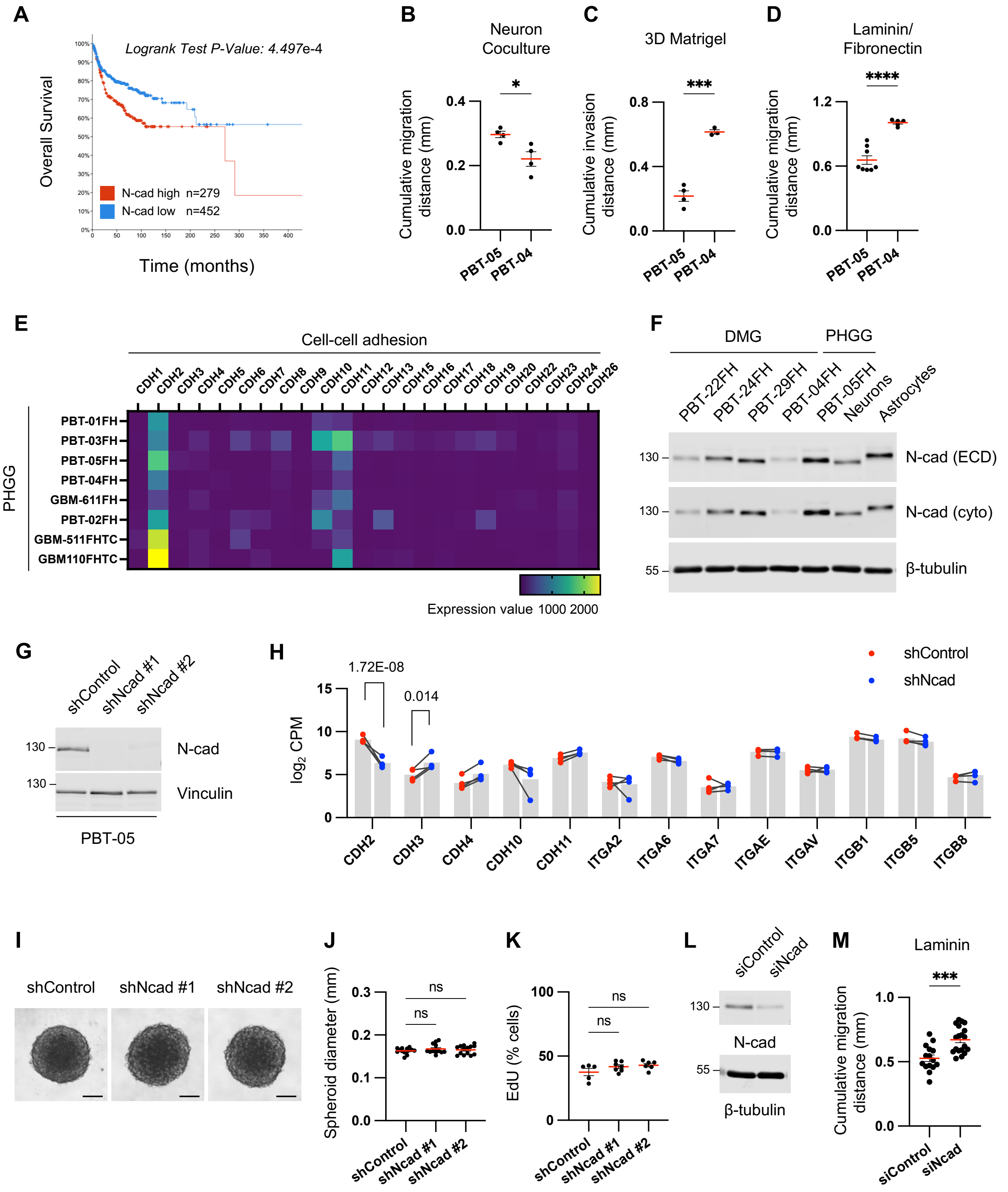
A
Logrank Test P-Value: 4.497e-4
Overall Survival
N-cad high n=279
N-cad low n=452
Time (months)
B
Neuron Coculture
Cumulative migration distance (mm)
PBT-05
PBT-04
C
3D Matrigel
Cumulative invasion distance (mm)
D
Laminin/ Fibronectin
Cumulative migration distance (mm)
E
Cell-cell adhesion
PHGG
PBT-01FH
PBT-03FH
PBT-05FH
PBT-04FH
GBM-611FH
PBT-02FH
GBM-511FHTC
GBM110FHTC
Expression value 1000 2000
F
DMG
PHGG
PBT-22FH
PBT-24FH
PBT-29FH
PBT-04FH
PBT-05FH
Neurons
Astrocytes
N-cad (ECD)
N-cad (cyto)
β-tubulin
G
shControl
shNcad #1
shNcad #2
N-cad
Vinculin
PBT-05
H
1.72E-08
0.014
shControl
shNcad
log2 CPM
CDH2
CDH3
CDH4
CDH10
CDH11
ITGA2
ITGA6
ITGA7
ITGAE
ITGAV
ITGB1
ITGB5
ITGB8
I
shControl
shNcad #1
shNcad #2
J
Spheroid diameter (mm)
K
EdU (% cells)
L
siControl
siNcad
N-cad
β-tubulin
M
Laminin
Cumulative migration distance (mm)

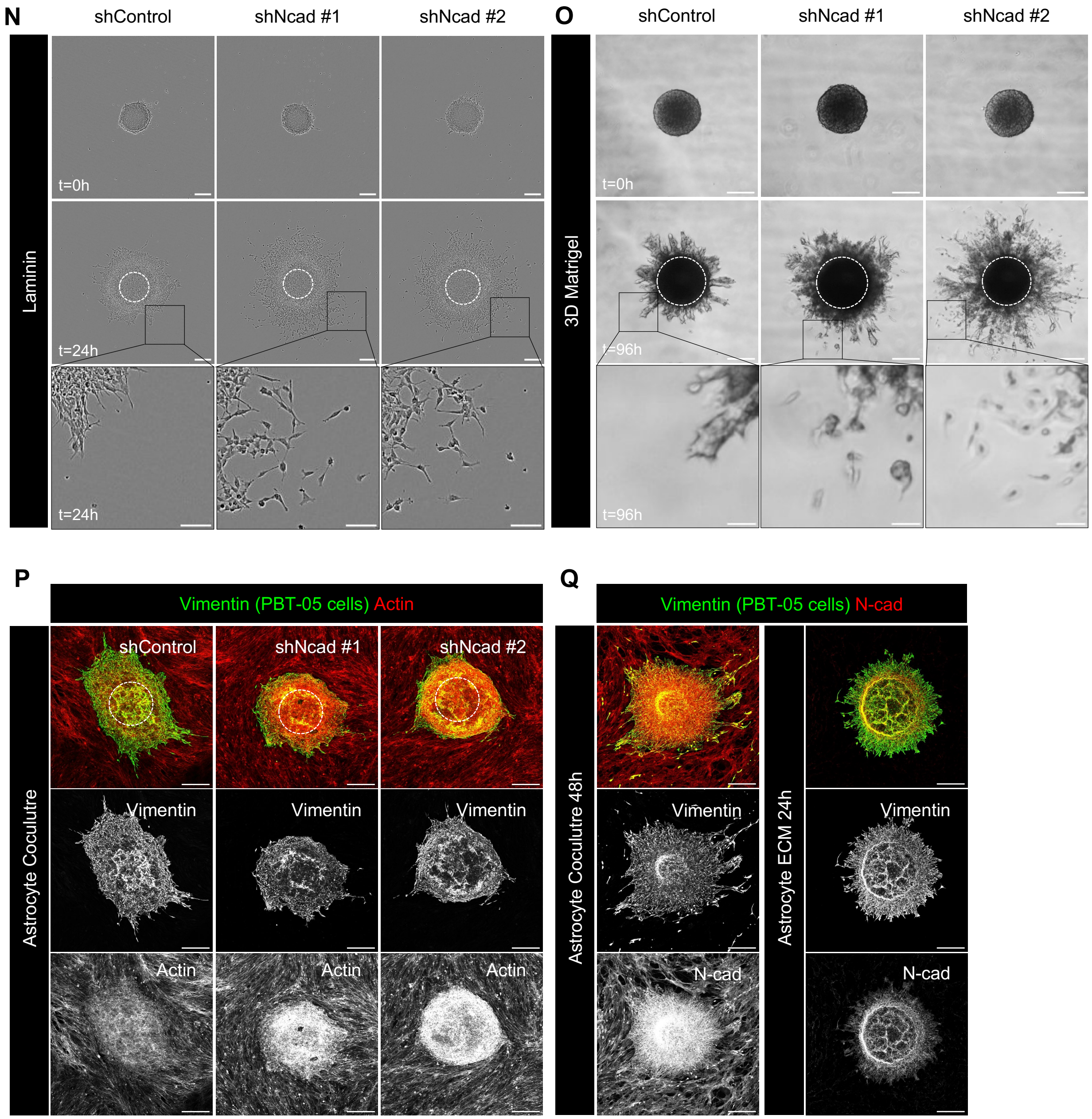


**Figure S1. N-cad expression, depletion, and controls for PHGG migration assays.** (A) Overall survival of pediatric brain tumor patients with high or low N-cad mRNA expression. Data collected from Pediatric Brain Tumor Atlas (PBTA) (http://pedcbioportal.kidsfirstdrc.org, PedcBioPortal, Open Pediatric Brain Tumor Atlas, v22). (B-D) Migration or invasion distance of PBT-04 and PBT-05 cells in mouse cerebellar neurons, Matrigel, and laminin/fibronectin. Error bars indicated mean$\pm$s.e.m. Unpaired *t*-tests. **P*<0.05, ****P*<0.001, *****P*<0.0001. (E) mRNA expression levels of cadherins in patient-derived PHGG tissues and cells (https://r2.amc.nl, R2: Genomics Analysis and Visualization, Mixed Pediatric PDX-Olson). (F) N-cad protein expression level in DMG and PHGG cell lines and mouse cerebellar neurons and mouse astrocytes. β-tubulin is shown as a loading control. (G) Western blots for control or N-cad shRNAs in PBT-05 cells. Vinculin is shown as a loading control. (H) RNA levels of Ncad and other cell adhesion proteins in shNcad cells. Mean CPM (counts per million) from 4 biological replicates. Lines connect paired samples. Adjusted *P* values from the Wald test using the Benjaminin-Hochberg method. (I-K) N-cad does not regulate spheroid formation or cell proliferation. (I) Representative images of spheroids. Scale bar, 100 μm. (J) Spheroid diameter at start of migration experiment. Error bars indicate mean±s.e.m. Ordinary one-way ANOVA Holm-Šídák's multiple comparisons test. ns, not significant. N=13-14 spheroids, 4 experiments. (K) EdU pulse labeling. Error bars indicate mean±s.e.m. Ordinary one-way ANOVA Holm-Šídák's multiple comparisons test. ns, not significant. N=5-7 spheroids, 3 experiments. (L) Western blots and (M) migration distance on laminin of control and N-cad siRNA-treated PBT-05 cells. Error bars indicate mean±s.e.m. Unpaired *t*-test. ***$P$<0.001. N=15-19 spheroids, 4 experiments. (N-O) N-cad depletion increases single-cell migration. Representative images of migrating control or N-cad shRNA spheroids on laminin or 3D Matrigel. Scale bars, 200 μm or 50 μm (inset) (P-Q) Controls for migration on astrocytes and astrocyte ECM. (P) Cell migration on mouse astrocytes. PBT-05 cells were identified with human-specific anti-vimentin antibodies. PBT-05 cell and astrocyte actin was detected with phalloidin. (Q) Decellularized astrocyte cultures lack N-cad. Scale bars, 200 μm.

### Figure S2

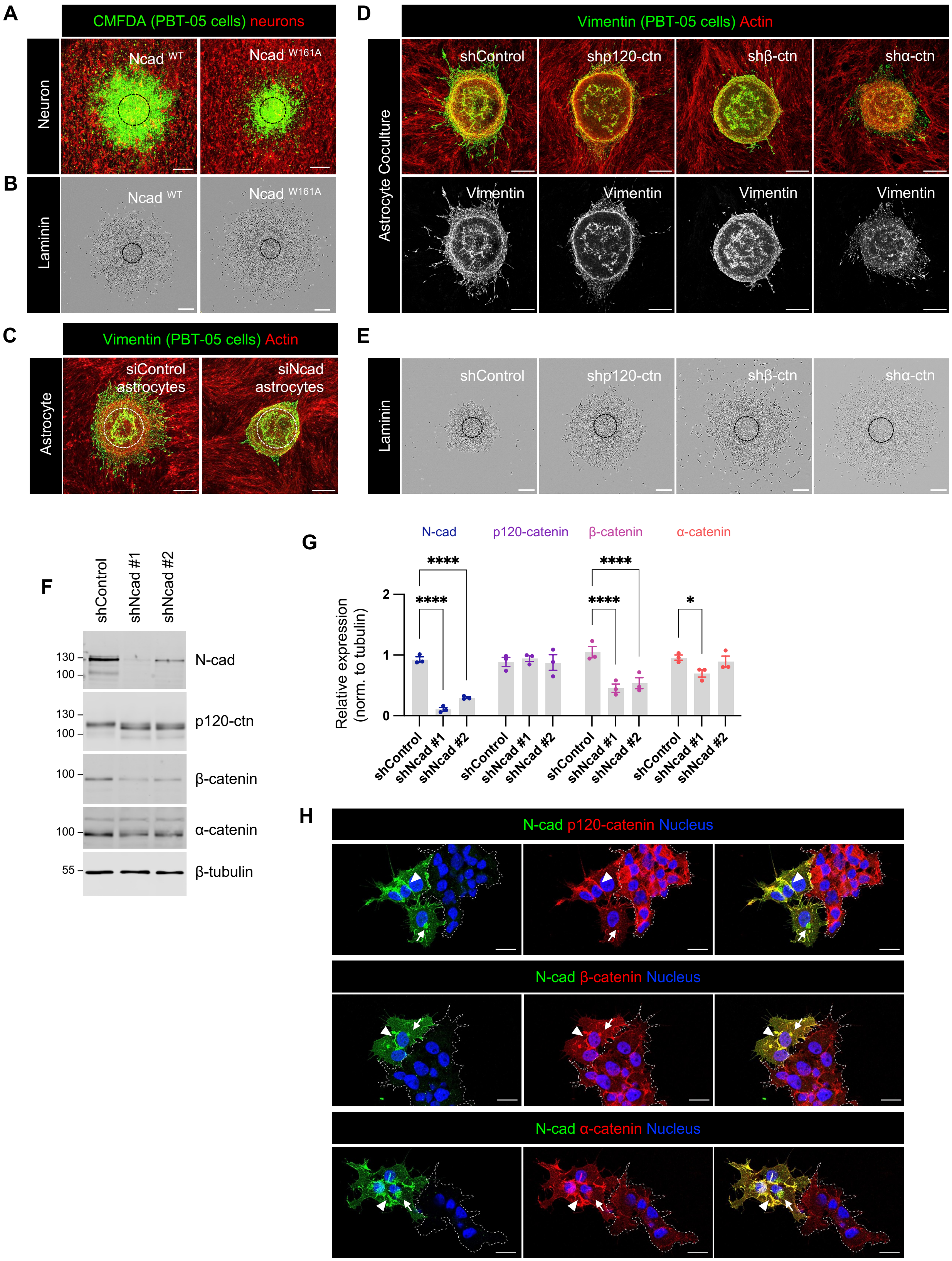

A
CMFDA (PBT-05 cells) neurons
Neuron
Ncad WT
Ncad W161A
B
Laminin
Ncad WT
Ncad W161A
C
Vimentin (PBT-05 cells) Actin
Astrocyte
siControl astrocytes
siNcad astrocytes
D
Vimentin (PBT-05 cells) Actin
Astrocyte Coculture
shControl
shp120-ctn
shβ-ctn
shα-ctn
Vimentin
E
Laminin
shControl
shp120-ctn
shβ-ctn
shα-ctn
F
shControl
shNcad #1
shNcad #2
130
100
N-cad
p120-ctn
β-catenin
α-catenin
55
β-tubulin
G
N-cad
p120-catenin
β-catenin
α-catenin
Relative expression (norm. to tubulin)
****
*
shControl
shNcad #1
shNcad #2
H
N-cad p120-catenin Nucleus
N-cad β-catenin Nucleus
N-cad α-catenin Nucleus

**Figure S2. Role of intercellular N-cad homotypic interactions and importance of N-cad for catenin localization.** (A-B) Representative images of $Ncad^{WT}$ and $Ncad^{W161A}$ expressing PBT-05 cells migrating on (A) cerebellar neurons or (B) laminin. (C) Representative images of PBT-05 cells migrating on siControl or siNcad-treated mouse astrocytes. Scale bars, 200 μm. (D-E) Images of PBT-05 cells expressing control, p120-catenin (p120-ctn), β-catenin (β-ctn) or α-catenin (α-ctn) shRNAs, migrating on (D) mouse astrocytes or (E) laminin. Scale bars, 200 μm. (F-G) N-cad, p120-catenin β-catenin and α-catenin protein levels in control and N-cad depleted cells. Protein expression levels were normalized to β-tubulin as a loading control. Two-way ANOVA Uncorrected Fisher's LSD test. N=3 Western blots. **P*<0.05, *****P*<0.0001. (H) Localization of p120-catenin, β-catenin and α-catenin in control or N-cad-depleted cells. Dashed lines indicate clusters of N-cad depleted cells. Arrowheads indicate cell-cell junctions and arrows indicate intracellular vesicles. Scale bars, 20 μm

### Figure S3

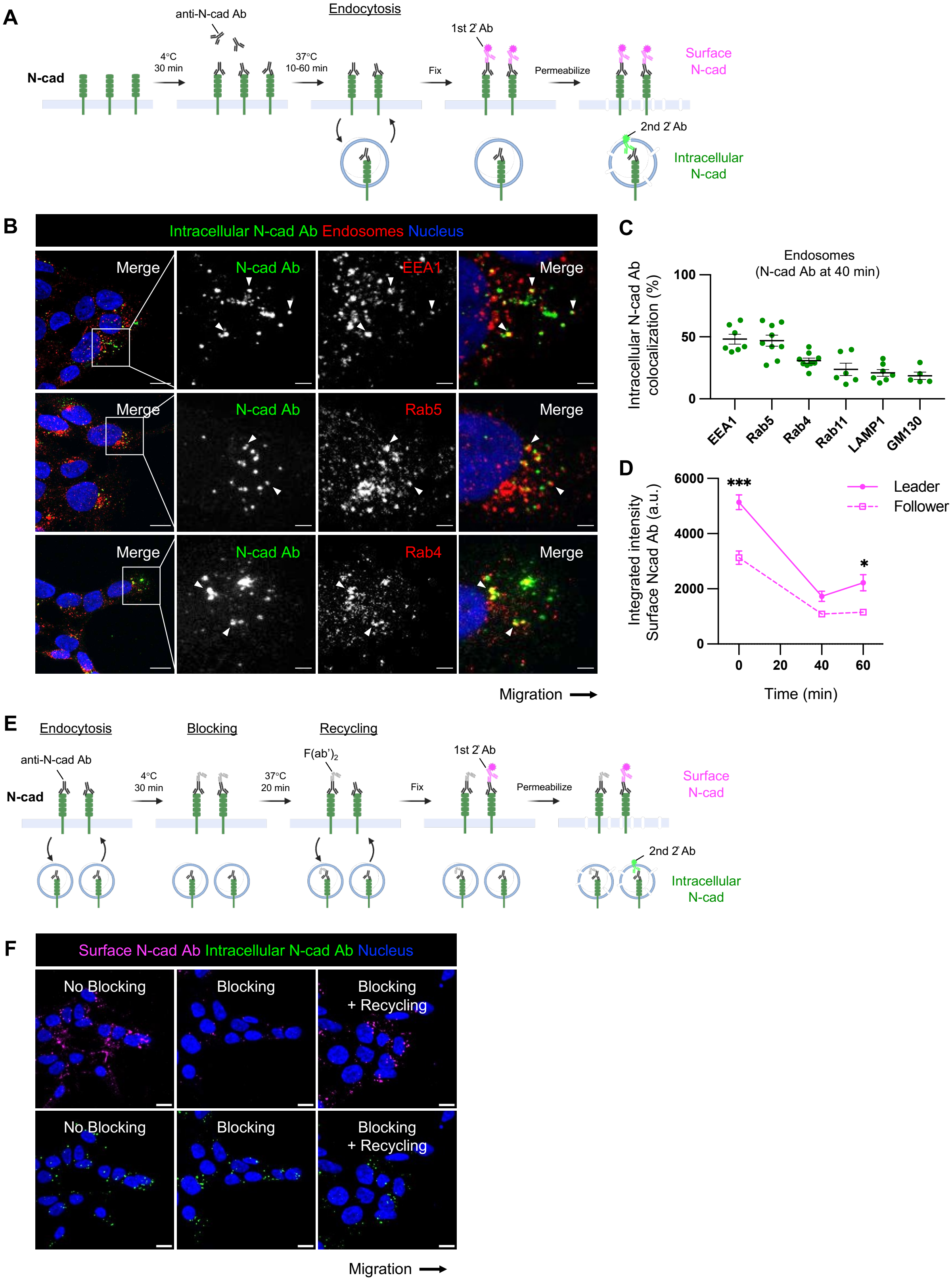
A
anti-N-cad Ab
Endocytosis
1st 2' Ab
N-cad
4°C
30 min
37°C
10-60 min
Fix
Permeabilize
Surface
N-cad
2nd 2' Ab
Intracellular
N-cad
B
Intracellular N-cad Ab Endosomes Nucleus
Merge
N-cad Ab
EEA1
Merge
Merge
N-cad Ab
Rab5
Merge
Merge
N-cad Ab
Rab4
Merge
Migration
C
Endosomes
(N-cad Ab at 40 min)
Intracellular N-cad Ab
colocalization (%)
100
50
0
EEA1
Rab5
Rab4
Rab11
LAMP1
GM130
D
***
*
Leader
Follower
Integrated intensity
Surface Ncad Ab (a.u.)
6000
4000
2000
0
0
20
40
60
Time (min)
E
Endocytosis
Blocking
Recycling
anti-N-cad Ab
N-cad
4°C
30 min
F(ab')2
37°C
20 min
1st 2' Ab
Fix
Permeabilize
Surface
N-cad
2nd 2' Ab
Intracellular
N-cad
F
Surface N-cad Ab Intracellular N-cad Ab Nucleus
No Blocking
Blocking
Blocking
+ Recycling
No Blocking
Blocking
Blocking
+ Recycling
Migration

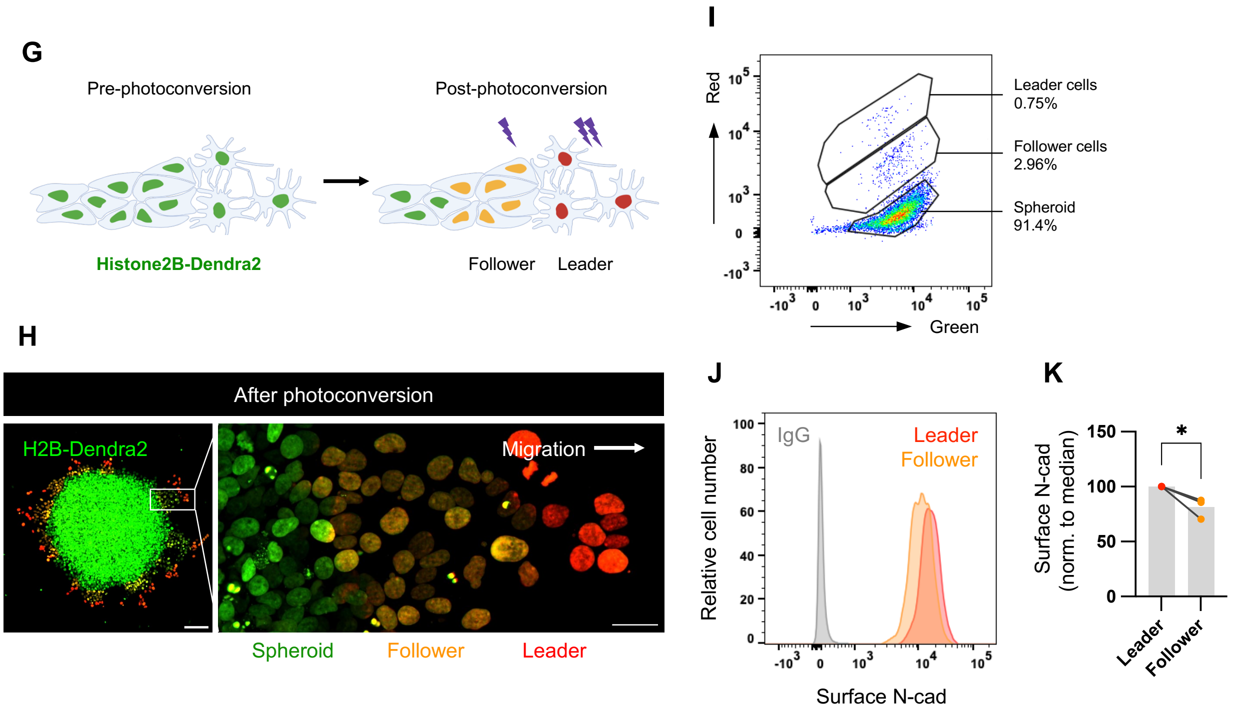


**Figure S3. N-cad endocytosis, recycling and surface levels in leader and follower cells.** (A) Schematic diagram of the N-cad antibody (Ab) internalization assay. Cells were labeled with N-cad ECD Ab in the cold, warmed for various times to allow internalization, fixed, and surface and internalized Ab detected with different fluorescent secondaries. (B) Immunofluorescence detection of internalized N-cad Ab with EEA1, Rab5 or Rab4 in leader cells after 40 min internalization. Arrowheads indicate colocalization. Scale bars, 10 $\mu$m or 2 $\mu$m (inset). (C) Object-based colocalization quantification of internalized N-cad Ab with EEA1, Rab5, Rab4, Rab11, LAMP1 or GM130. N=5-10 spheroids, 3 experiments. (D) Surface N-cad Ab increases between 40 and 60 min. a.u., arbitrary units. Two-way ANOVA Šídák's multiple comparisons test multiple comparisons test. *$P$<0.05, ***$P$<0.001. N=3-4 spheroids, 3 experiments. (E-F) N-cad Ab recycling assay. (E-F) Schematic and representative images of N-cad recycling assay. Cells were incubated with N-cad ECD antibody in the cold and warmed for 40 min to allow internalization. N-cad Ab remaining on the surface was blocked in the cold with excess $F(ab')_2$. Cells were then warmed to allow recycling before fixation and detection of surface and internalized Ab with different fluorescent secondaries. Scale bars, 10 $\mu$m. (G) Schematic diagram and (H) representative images showing photoconversion of histone H2B-Dendra2 in leader and follower cells. (I-K) Flow cytometry. (I) Definition of leader, follower and spheroid cells according to extent of photoconversion. (J) N-cad surface intensity histograms. (K) Normalized median N-cad fluorescent intensity. Paired *t*-test. *$P$=0.0374. N=3 experiments.

### Figure S4

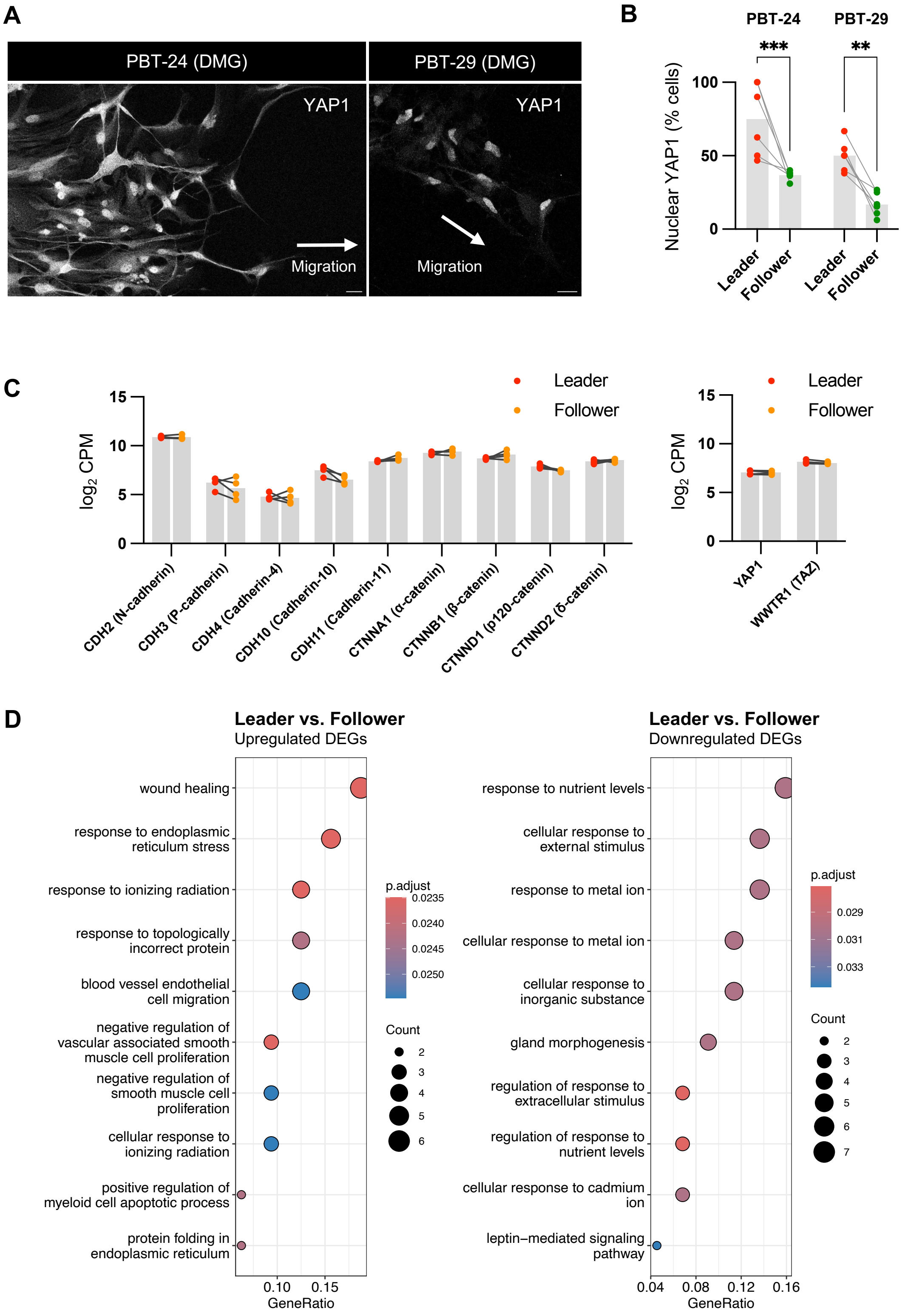


**Figure S4. YAP1 signaling and wound healing gene expression is increased in leader cells.** (A-B) Representative images and quantification of YAP1 localization in leader (L) and follower (F) cells in PBT-24 and PBT-29 DMG cells migrating on Matrigel for 72 hrs. Scale bars, 20 μm. N=5 spheroids, 3 experiments. Two-way ANOVA Šídák's multiple comparisons test. ***P*<0.01, ****P*<0.001. (C) RNASeq analysis of cadherin, catenin, YAP1 and TAZ expression in leader and follower cells. (D) Gene ontology functional enrichment analysis of differentially-expressed genes (DEGs) between leader and follower cells.

### Figure S5

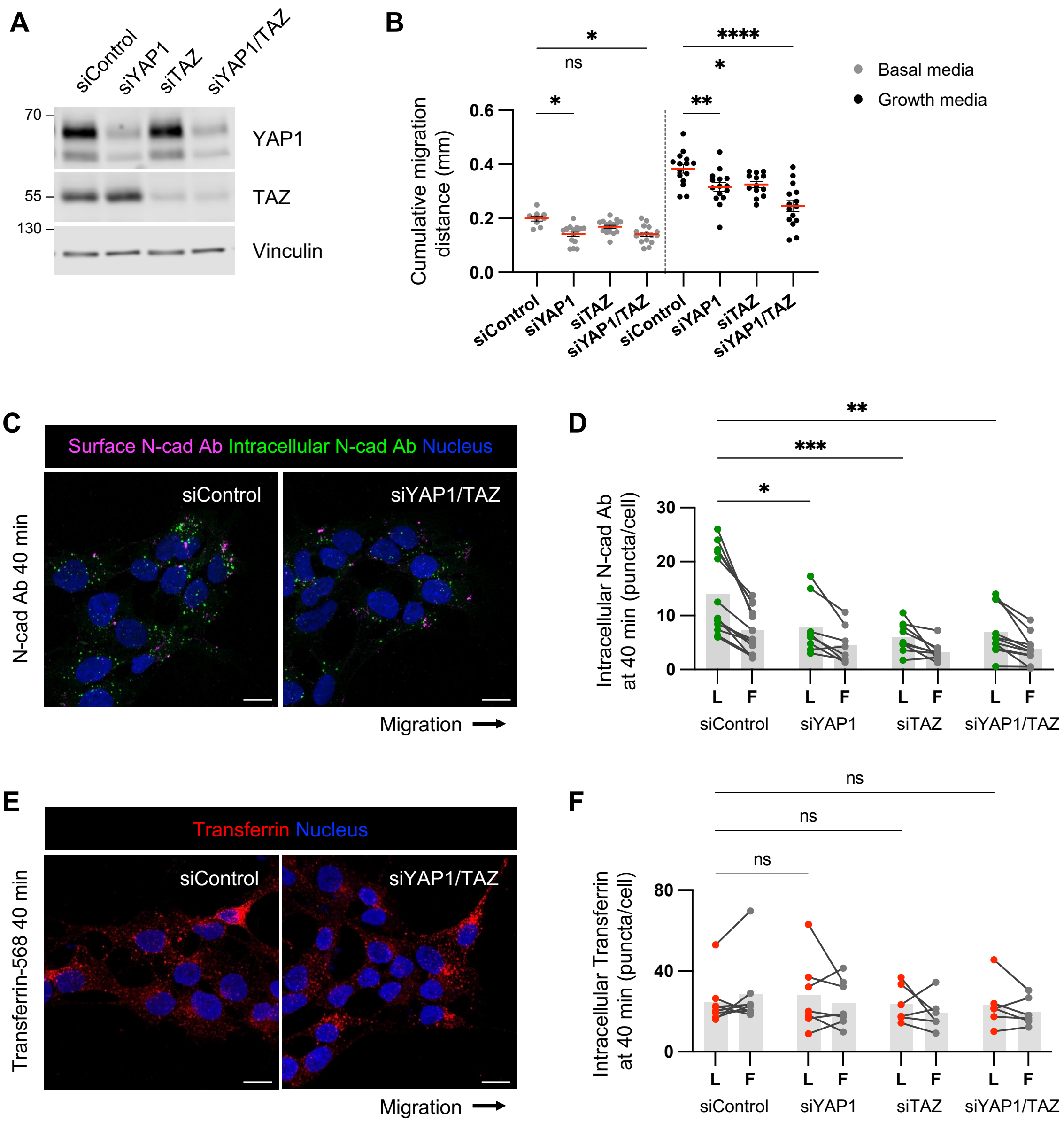


**Figure S5. YAP1/TAZ regulates PHGG migration and N-cad endocytosis.** (A) Western blot analysis of control, YAP1, TAZ or YAP1/TAZ siRNAs in PBT-05 cells. Vinculin is shown as a loading control. (B) Cumulative migration distance on laminin for 24 hrs. Ordinary One-way ANOVA Šídák's multiple comparisons test. N=9-16 spheroids, 3 experiments. (C-D) Representative images and quantification of internalized N-cad Abs at 40 min incubation. N= 4-13 spheroids, 3 experiments. (E-F) Representative images and quantification of internalized Alexa Fluor 568-conjucated transferrin at 40 min incubation. N= 5-7 spheroids, 2 experiments.

(D, F) Two-way ANOVA Šídák's multiple comparisons test. $^*P<0.05$, $^{**}P<0.001$, $^{***}P<0.001$, $^{****}P<0.0001$. ns, not significant.

**Table S1.** Cell adhesion receptors in PHGGs. mRNA expression levels of cell adhesion receptors in patient-derived PHGG tissues and cells (https://r2.amc.nl, R2: Genomics Analysis and Visualization, Mixed Pediatric PDX-Olson).

**Table S2.** RNA sequencing results comparing control and N-cad shRNA cells.

**Table S3.** RNA sequencing results comparing leader and follower cells.

**Video 1.** N-cad inhibits PHGG cell migration and cell dissociation. Control or N-cad shRNA PBT-05 spheroids were imaged every 15 min for 48 hrs during migration on laminin. Images were captured by Incucyte, Scale bars, 200 μm.

**Video 2.** Leader and follower cells interconvert during PHGG migration. Histone H2B-Dendra2 expressing PBT-05 cells migrating on neurons or laminin. Time-lapse images were taken every 15 min for 16 hrs. Images were captured by spinning disk confocal microscope. Scale bars, 20 μm.
